# OJIP chlorophyll fluorescence induction profiles and plastoquinone binding affinity of the Photosystem II assembly intermediate PSII-I from *Thermosynechococcus elongatus*

**DOI:** 10.1101/2021.06.28.450235

**Authors:** Jure Zabret, Marc M. Nowaczyk

## Abstract

Binding of Psb28 to the photosystem II assembly intermediate PSII-I induces conformational changes to the PSII acceptor side that impact charge recombination and reduce the *in situ* production of singlet oxygen (Zabret et al. 2021, Nat. Plants 7, 524-538). A detailed fluorometric analysis of the PSII-I assembly intermediate compared with OEC-disrupted and Mn-depleted PSII complexes showed differences between their variable (OJIP) chlorophyll fluorescence induction profiles. These revealed a distinct destabilisation of the Q_A_^-^ state in the PSII-I assembly intermediate and inactivated PSII samples related to an increased rate of direct and safe charge recombination. Furthermore, inactivation or removal of the OEC increases the binding affinity for plastoquinone analogues like DCBQ to the different PSII complexes. These results might indicate a mechanism that further contributes to the protection of PSII during biogenesis or repair.

## 1. Introduction

Oxygenic photosynthesis, the conversion of light energy into biologically usable chemical energy, is the primary reaction needed to sustain life on Earth. It forms the basis of our food chain by converting atmospheric CO_2_ into biomass, but it also dictates our atmosphere’s balance by splitting water and releasing molecular oxygen. The water-splitting and oxygen-evolving reactions are carried out by photosystem II (PSII), a multi-subunit homodimeric transmembrane protein complex found in thylakoid membranes of cyanobacteria and chloroplasts of algae and higher plants [2,3].

In PSII, absorbed light energy is channelled to the reaction center chlorophylls (P_680_). After the initial charge separation, an electron is released and transferred via pheophytin and the immobile plastoquinone (Q_A_) to mobile plastoquinone (Q_B_). It in turn leaves the binding pocket in the form of hydroquinone (PQH_2_) after accepting two electrons from Q_A_ and two protons from D1 His252 and D1 His215, respectively [4]. The open binding site is occupied again with an oxidized plastoquinone molecule from the plastoquinone pool within the thylakoid membrane. The oxidized reaction center (P_680_^+^) is re-reduced by a nearby tyrosine residue (Tyr_Z_) that transfers electrons from the oxygen-evolving cluster (OEC), which catalyzes water oxidation [5].

Each PSII monomer comprises up to 20 protein subunits, 13 of which are classified as low molecular weight (<10 kDa) subunits, 2-3 plastoquinone, two pheophytin, two heme, 35 chlorophyll and 12 carotenoid molecules, one none-heme iron, and the unique Mn_4_CaO_5_ cluster responsible for water oxidation. Additionally, PSII contains three Cl^-^ and 20 to 25 lipid molecules [6], with most components performing a distinct function. Assembly (or biogenesis) of this large protein complex is spatially and temporally organized in all organisms. It involves an assembly-line like process facilitated by numerous auxiliary protein factors. These are characterized by their transient interaction with PSII [7–9].

The auxiliary protein Psb28, the only cytoplasmatic PSII auxiliary protein, found in cyanobacteria and higher plants [10], is involved in PSII biogenesis and repair [11]. It interacts with the RC47 module. More specifically it binds to the cytoplasmatic side of the CP47 subunit [12]. Furthermore, deletion of Psb28 in *Synechocystis* sp. PCC 6803 showed a strong photosynthetic phenotype. This was apparent in the slower autotrophic growth of the mutant [11,12]. Moreover, Psb28 deletion strains exhibit a slower PSII recovery rate after light induced photodamage [11] and an overall increased PSII turnover rate [12].

Recently, we confirmed that the auxiliary factor Psb28 plays a protective role in the PSII-I assembly intermediate [1]. By reorienting the D-E loop of the D1 subunit, Psb28 changes the position of D1 Phe265, Ser264 and Tyr252. This conformational change disrupts the Q_B_-binding pocket and prevents forward electron transfer across the PSII acceptor side. Reorientation of the D-E loop also causes the sidechain of D2 Glu241 to bind to the non-heme iron in place of the bicarbonate ligand [1] (Fig. 1). Removal of the bicarbonate from PSII induces a positive shift in Q_A_/Q_A_^-^ redox potential, favoring direct recombination between the P_680_^+^/Q_A_^-^ charge pair, thereby bypassing the triplet chlorophyll state and minimizing singlet oxygen (^1^O_2_) production [1,13], as measured in the PSII-I assembly intermediate [1]. This is needed since PSII is most prone to photoinactivation by reactive oxygen species (ROS) during the assembly process [14].

**Fig. 1:**
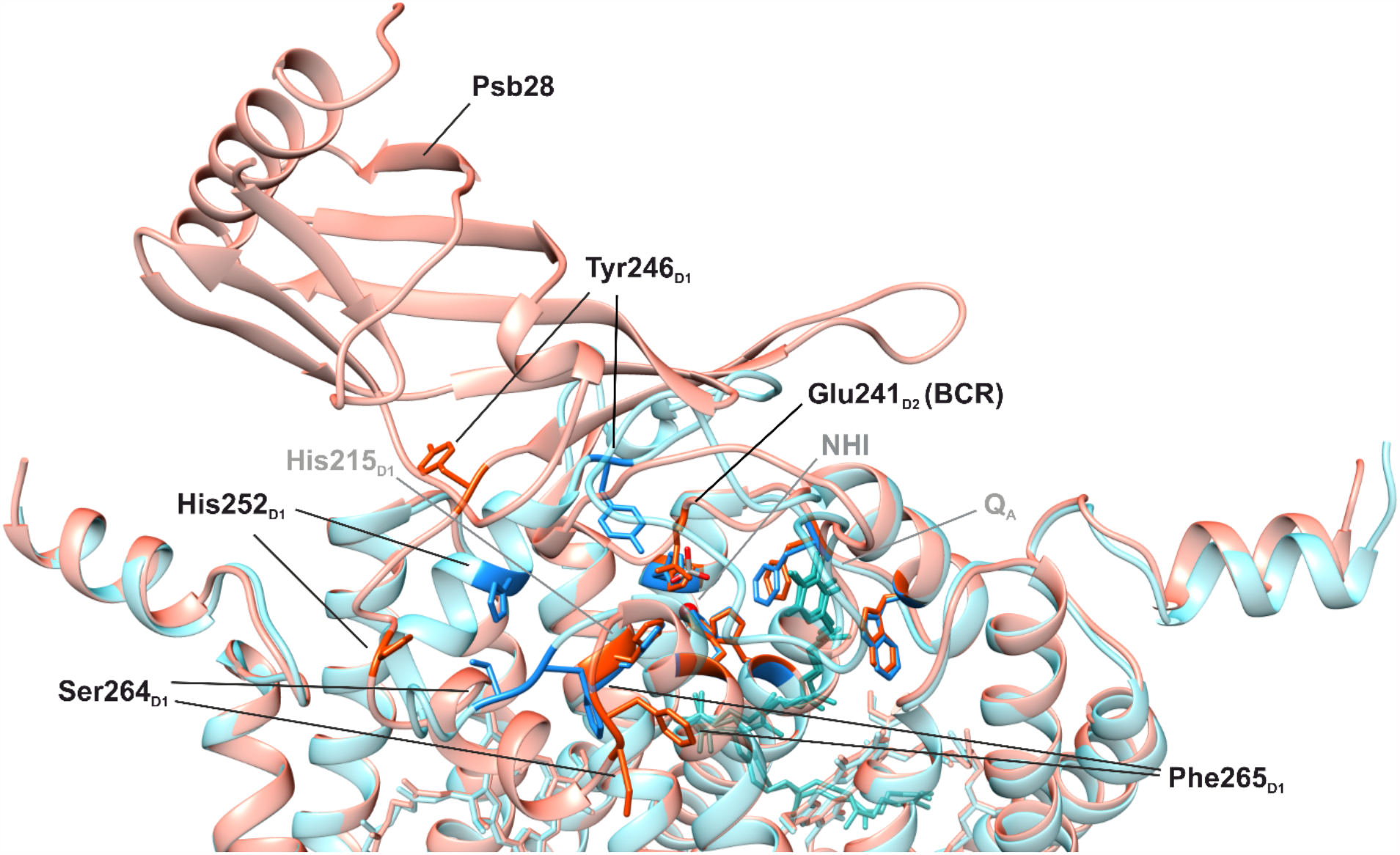
Distortion of the Q_B_-binding pocket caused by the Psb28 auxiliary protein. Structural alignment of the PSII-I (PDB ID: 7HNP) assembly intermediate (red) and the PSII-M monomer (PDB ID: 7NHO) (blue) core subunits D1 and D2, created using UCSF Chimera [15]. Residues and structures marked in grey are not altered, and the binding of Psb28 alters the residues marked in black.

In this paper, we use the highly purified PSII-I assembly intermediate [1], CaCl_2_-washed [16], and Mn-depleted PSII [17] in comparison with active PSII preparations to gauge the effect of OEC disruption and removal on Q_A_ reduction and oxidation kinetics trough alterations in their variable (OJIP) chlorophyll fluorescence induction profiles. Furthermore, we quantified the interaction between the different PSII complexes and the artificial electron acceptor 2,5-dichloro-1,4-benzoquinone (DCBQ). We show that inactive PSII complexes have a higher affinity towards DCBQ than active PSII complexes.

## 2. Methods

### 2.1. Culture conditions

*T. elongatus* CP43-TS and *T. elongatus* ΔpsbJ CP43-TS cells were grown as described previously. In brief, cultures were cultivated in BG-11 liquid medium using 25-litre foil fermenters (Bioengineering) at 45 °C, 5 % (v/v) CO_2_-enriched air bubbling and 50-200 µE white light illumination. Cells were harvested at an OD_680_ of approximately two and concentrated using an Amicon DC10 LA hollow-fiber system. This was followed by centrifugation (3500 rcf and 25 °C for 45 min), and lastly, the cells were resuspended in buffer D (100 mM Tris-HCl, pH 7.5, 10 mM MgCl_2_, 10 mM CaCl_2_, 500 mM mannitol and 20% (w/v) glycerol), flash-frozen in liquid N_2_ and stored at -80 °C until further use.

### 2.2. Isolation and PAGE analysis of Strep tagged PSII

Cell disruption and thylakoid membrane preparation, followed by Strep-Tactin-affinity purification of CP43-tagged PSII complexes, separated via anion exchange chromatography and subsequent PAGE analysis (Blue-native and SDS-PAGE) were performed as described in our previous work [1].

### 2.3. CaCl_2_ washing and oxygen evolution assays with isolated PSII complexes

The extrinsic PSII subunits PsbO, PsbU and PsbV were removed by applying active dimeric PSII onto a size exclusion column (Superdex 75 10/300 GL, GE Healthcare) pre-equilibrated in CaCl_2_ buffer (1 M CaCl2, 10 mM MgCl_2_, 20 mM MES-NaOH, pH 6.5, 0.03% (w/v) DDM) [1,16]. OEC-disrupted PSII samples (PSII(-OUV)) were collected and washed in activity buffer (150 mM KCl, 20 mM MES-KOH, pH 6.5, 10 mM MgCl_2_, 10 mM CaCl_2_ and 0.03% (w/v) DDM) using a centrifugal concentrator unit (Amicon, Ultra – 15, 100 000 NMWL).

### 2.4. Removal of the PSII oxygen-evolving cluster

The Mn-cluster was removed from isolated dimeric PSII as described by [17]. In brief, PSII complexes were diluted to a chlorophyll concentration of 0.1 mg/ml in Mn-depletion buffer (20 mM MES-NaOH, pH 6.5, 10 mM MgCl_2_, 10 mM CaCl_2_ and 500 mM mannitol, 0.03 % (w/v) DDM, 10 mM NH_2_OH and 1 mM EDTA) and incubated at 4 °C for 60 min. The resulting apo-PSII sample was equilibrated in activity buffer (100 mM KCl, 20 mM MES-KOH, pH 6.5, 10 mM MgCl_2_, 10 mM CaCl_2_ and 0.03% (w/v) DDM) using a spin concentrator (Amicon, Ultra – 15, 100 000 NMWL).

### 2.5. PSII oxygen evolution rates

Light-induced PSII oxygen evolution rates, with ferricyanide (FeCy) and 2,6-dichloro-p-benzoquinone (DCBQ) as artificial electron acceptors, were performed as described previously [18].

### 2.6. Variable chlorophyll fluorescence induction

Isolated PSII samples were diluted in activity buffer (150 mM KCl, 20 mM MES-KOH, pH 6.5, 10 mM MgCl_2_, 10 mM CaCl_2_ and 0.03% (w/v) DDM) to a final reaction centre concentration of 150 nM in a volume of 3 ml and dark-adapted for 5 min before measurements. Chlorophyll fluorescence induction traces were collected using an FL3500 Dual-Modulation Kinetic fluorometer (PSI, Photon Systems Instruments). Red actinic light power was set to 80 %, and traces were collected for 120 seconds in three successive measurements. When present, the concentration of 3-(3,4-dichlorophenyl)-1,1-dimethylurea (DCMU) or 2,6-dichloro-p-benzoquinone (DCBQ) were 20 µM. Variable chlorophyll fluorescence induction traces obtained without the addition of DCMU and traces obtained with the PSII assembly intermediate PSII-I in the presence of DCMU were deconvoluted using the sum of three first-order kinetics by nonlinear regression using Sigma Plot [19]. PSII and PSII(-OUV), in the presence of DCMU, exhibited fluorescence induction traces which were best fitted using a single first-order equation;

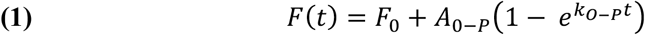

where *F(t)* is fluorescence at time *t, F*_*0*_ being the initial fluorescence, *A*_*O-P*_ the amplitudes, and *k*_*O-P*_ the rate constants of the single fluorescence transient.

### 2.7. DCBQ titration assays

The binding affinity of DCBQ to various PSII complexes was determined via titration of the quenching effect of DCBQ on chlorophyll fluorescence. A 16-point serial dilution (1:1) of DCBQ was prepared in titration buffer (150 mM KCl, 20 mM MES-KOH, pH 6.5, 10 mM MgCl_2_, 10 mM CaCl_2_, 0.03% (w/v) DDM and 5 % (v/v) methanol), with the concentration ranging from 5 mM to 153 nM. The concentration of PSII reaction centres was 900 nM throughout the dilution series. Chlorophyll fluorescence values were recorded with a Monolith NT 115 system (Nano Temper Technologies), using 5 % red LED power and Standard capillaries (Nano Temper Technologies) at room temperature. Initial fluorescence data was evaluated using the Graph Pad Prism 8.0 software, using the specific binding with Hill slope equation;

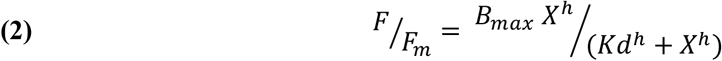

where *B*_*max*_ is the maximum specific binding, *Kd* the equilibrium dissociation constant and *h* the Hill slope.

## 3. Results

### 3.1. Combination of Strep-tag-affinity purification and anion exchange chromatography enabled isolation of PSII complexes at various assembly stages

The fusion of a Twin-Strep-tag (TS) with the C-terminus of the CP43 subunit in *T. elongatus* and *T. elongatus* ΔpsbJ [1] enabled the efficient purification of several different PSII complexes. First, proteins were isolated via Strep-tag-affinity chromatography and separated into different PSII subfractions with anion exchange chromatography. Next, the oligomeric states and subunit composition of active PSII and the PSII-I assembly intermediate were examined using BN-PAGE and SDS-PAGE, respectively (Fig. 2a and b).

**Fig. 2:**
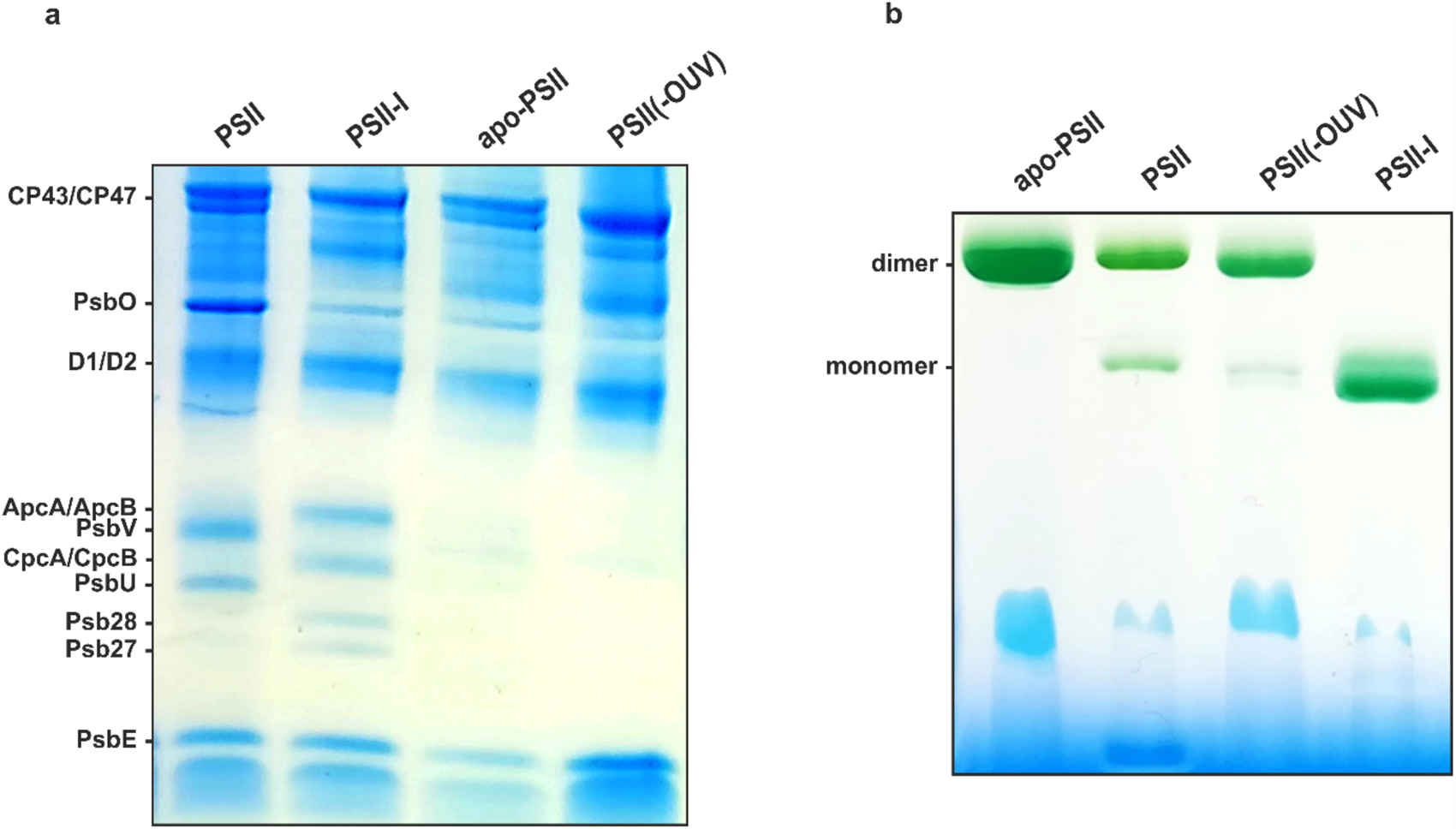
PAGE analysis of purified PSII complexes. **(a)** SJ-PAGE and **(b)** BN-PAGE of the Psb27, Psb28 and Psb34 containing PSII assembly intermediate (PSII-I), active dimeric PSII (PSII), CaCl_2_-washed PSII (PSII(-OUV)) and hydroxylamine-treated PSII (apo-PSII).

A detailed biochemical characterisation of the isolated core complexes (PSII and PSII-I) can be found elsewhere [1,18,20]. In brief, a monomeric PSII assembly intermediate in complex with the auxiliary proteins Psb27, Psb28 and Psb34, termed PSII-I, was isolated from *T. elongatus* CP43-TS where the low molecular weight (LMW) subunit PsbJ was deleted through insertional inactivation of its coding sequence [1,20]. In addition, active dimeric PSII (PSII) was isolated from *T. elongatus* CP43-TS [1]. The isolated active dimeric PSII fraction exhibited an oxygen-evolving activity of 2492 ± 286 µmol O_2_ (mg_Chl_ h)^-1^. The same protein fraction was used to prepare OEC disrupted dimeric (PSII(-OUV)) and Mn-depleted PSII (apo-PSII) samples.

As reported previously [1,16], CaCl_2_-washing led to the disassociation of the extrinsic subunits PsbO, PsbU and PsbV. We saw a drastic decrease in the band intensity for all extrinsic subunits, indicating an almost complete disruption of the OEC (Fig. 2a, PSII(-OUV)). However, PsbO is still present in sub-stoichiometric amounts. The oxygen-evolving activity of CaCl_2_-washed PSII (PSII(-OUV) decreased approximately 10-fold to 197 ± 14 µmol O_2_ (mg_Chl_ h)^-1^.

Removing the OEC by hydroxylamine treatment [17] was accompanied by disassociation of the extrinsic subunits PsbU and PsbV, with only traces of PsbO remaining attached to the complex (Fig. 2a, apo-PSII). Moreover, efficient Mn-cluster removal was apparent from a complete absence of oxygen-evolving activity of the apo-PSII sample.

On the other hand, the PSII-I assembly intermediate lacks the extrinsic subunits PsbU and PsbV, with PsbO present only in sub-stoichiometric amounts (Fig. 2a, PSII-I). Moreover, the assembly intermediate has a detectable density at the high-affinity site (HAS). Most likely, this is an Mn atom, but we could not exclude the presence of a different cation without further analysis [1]. The lack of an Mn-cluster is translated into a residual oxygen-evolving activity of less than 50 µmol O_2_ (mg_Chl_ h)^-1^ indicating slight, but negligible, contamination with active PSII.

### 3.2. Disruption or removal of the OEC leads to the onset of the co-called K-peak in the OJIP fluorescence transient

PSII fluorescence yield is determined by the oxidation state of Q_A_, the immobile plastoquinone. Fully open PSII reaction centres (RCs) are characterized by an oxidized Q_A_ and a low fluorescence yield accompanied by a high electron trapping efficiency (Φ_tr_). Conversely, when Q_A_^-^ is present, the RC is considered closed. The closed state is characterized by a comparably high fluorescence yield and a low Φ_tr_, which leads to a low probability of charge stabilization in the Q_A_^-^ state [21,22]. Fast variable (OJIP) chlorophyll fluorescence induction is determined by these characteristics. The fluorescence transient goes through four distinct phases (O, J, I and P). First, the fluorescence traces begin at the fluorescence origin (*F*_*0*_), indicated by the O-feature. This is followed by the J-peak, with the O-J rise reflecting the accumulation of reduced immobile plastoquinone (Q_A_^-^). From this, the fluorescence signal continues to rise through the intermediary inflexion (I). This last step is believed to represent the further closing of RCs by the accumulation of Q_A_^-^Q_B_ ^2-^ [23], yet others have connected this feature with events on the donor side of PSII [24]. After this, the traces reach the maximal fluorescence value (*Fm*) in the P-peak. At this point, the concentration of Q_A_^-^Q_B_ ^2-^ and PQH_2_ is considered to be maximal [23,25].

In our control PSII sample, the O-J rise accounts for 82.3 % of the fluorescence amplitude. The J-peak is followed by the J-I and I-P transitions, which account for 4.7 and 8.7 % of the signal, respectively. Removal of the Mn-cluster in the apo-PSII sample lowered the amplitude of the O-J rise to 39.1 % (a decrease of 43.2 %) and increased the J-I and I-P transition to 27.9 and 33.9 %, respectively. In contrast, OEC disruption only decreased the J-peak amplitude to 59.5 % (a decrease of 22.8 %). We can see a similarly low O-J transition amplitude of 36.6 % in the PSII-I assembly intermediate, with the J-I and I-P transition accounting for 22.7 and 40.5 % of the fluorescence increase, respectively (Table 1).

**Table 1:** Variable chlorophyll fluorescence induction best-fit kinetics values.

| | *Amp. (%)* | *PSII* | ± | *PSII(-OUV)* | ± | *apo-PSII* | ± | *PSII-I* | ± |
| --- | --- | --- | --- | --- | --- | --- | --- | --- | --- |
| **no addition** | *O-J* | 82.3 | 0.1 | 59.5 | 0.2 | 39.1 | 0.001 | 36.6 | 0.1 |
|  | *J-I* | 4.7 | 0.04 | 14.7 | 0.1 | 27.9 | 0.007 | 22.7 | 0.6 |
|  | *I-P* | 8.7 | 0.04 | 19.2 | 0.1 | 33.9 | 0.01 | 40.5 | 0.6 |
| **20 μM DCBQ** | *O-J* | 59 | 0.1 | 64 | 0.1 | 58.6 | 0.1 |  |  |
|  | *J-I* | 7.4 | 0.2 | 2.6 | 0.1 | 10.4 | 0.6 | n.a. | n.a. |
|  | *I-P* | 37 | 0.2 | 42 | 0.4 | 30.3 | 0.01 |  |  |
| **20 μM DCMU** | *O-J* |  |  |  |  | 53.1 | 0.0003 | 42 | 0.1 |
|  | *J-I* | n.a. | n.a. | n.a. | n.a. | 12.2 | 0.002 | 39 | 0.4 |
|  | *I-P* |  |  |  |  | 34.8 | 0.002 | 20 | 0.5 |

Similar changes to chlorophyll fluorescence induction profiles were observed in Mn-deficient barley (*Hordeum vulgare*) mutant lines [26]. The lack of a functional Mn-cluster altered the OJIP fluorescence induction profile by shifting the J-peak forward and lowering its intensity. This type of change in the OJIP fluorescence induction profile is called the K-peak and is connected with an inactive or absent Mn-cluster [26–29]. Its removal induces a positive shift in Q_A_/Q_A_^-^ redox potential, accompanied by slight structural perturbations that shift the primary quinone’s redox potential by approximately +150 mV [30]. Compared with the control PSII sample, we observed decreases in the O-J transition amplitudes in the artificially created apo-PSII and PSII(-OUV) samples and the isolated PSII-I assembly intermediate, which either lack an OEC or had it inactivated (Table 1 and Fig. 3a). This occurrence may indicate that the J-peak amplitude can be used to assess the Q_A_/Q_A_^-^ redox potential in isolated PSII complexes.

**Fig. 3:**
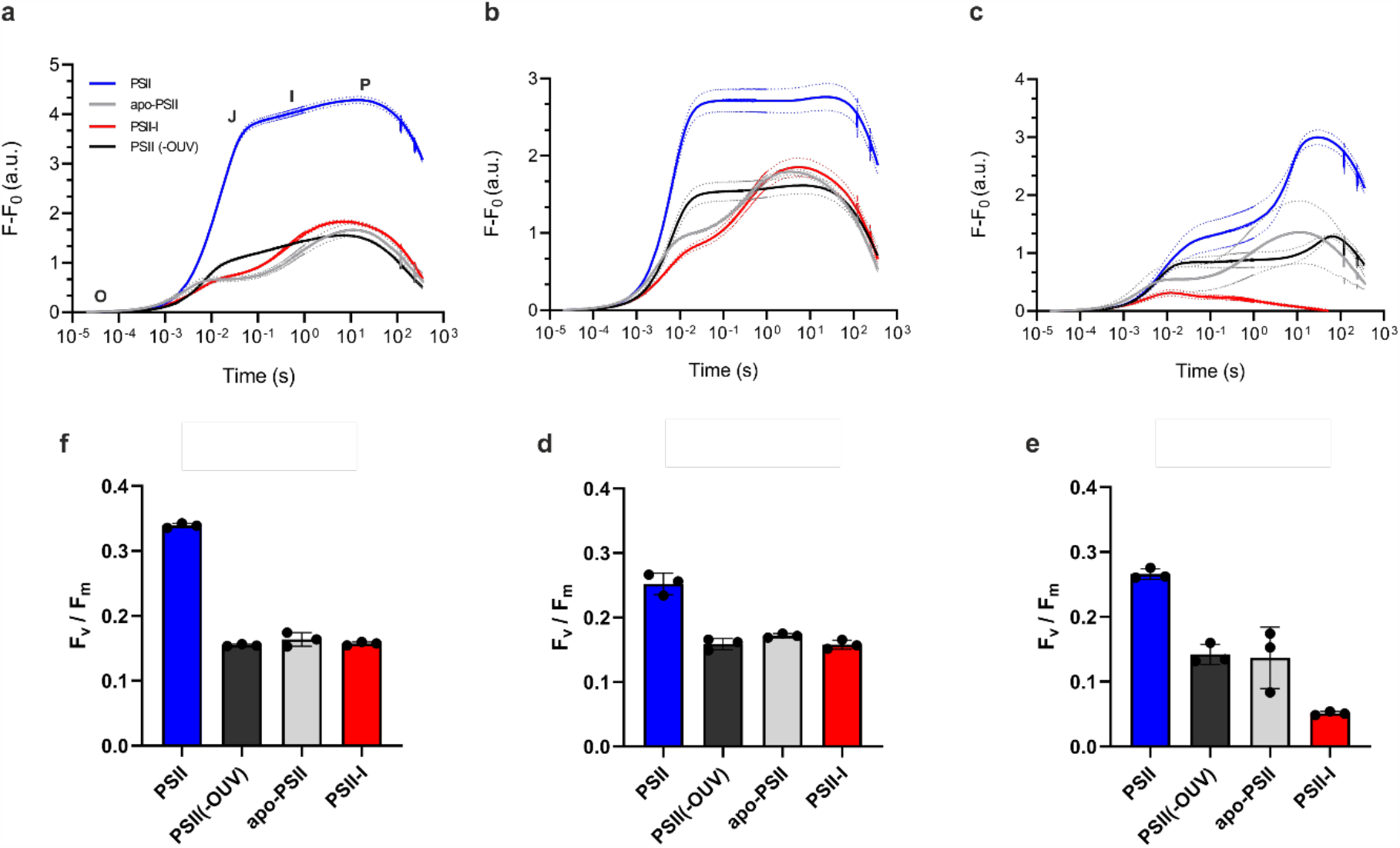
**Variable chlorophyll a fluorescence induction** with **(a)** no addition, **(b)** 20 µM DCMU, and **(c)** 20 µM DCBQ. Fluorescence traces were baseline corrected, and dotted corridors depict the standard deviation (SD) three independent measurements (*n* = 3). **Calculated Fv/Fm ratios with**, calculated form OJIP fluorescence induction traces, with **(d)** no addition, **(e)** 20 µM DCMU, and **(f)** 20 µM DCBQ. Error bars depict the standard deviation (SD) three independent measurements (*n* = 3).

### 3.3. Blocking forward electron flow increased the J-peak intensity in the OJIP fluorescence transient

Interestingly, the addition of DCMU has a similar effect on the control PSII sample as it does on the OEC disrupted PSII(-OUV) sample (Fig. 3b, blue and black traces). Both protein complexes exhibit a drastic increase in J-peak intensity, equal to or close to the maximum fluorescence value (*Fm*) in the P-peak. A similar J-peak increase after the addition of DCMU was observed with whole-cell *Synechocystis* sp. PCC 6803 variable chlorophyll fluorescence induction measurements [31,32]. This drastic increase in J-peak intensity indicates that blocked forward electron flow allows for a faster complete reduction of Q_A_, the primary electron acceptor [33]. On the other hand, the Mn-depleted apo-PSII sample showed a much smaller relative J-peak intensity increase (Fig. 3b, grey trace), similar to trends observed in hydroxylamine treated canola leaf discs [34]. With the addition of DCMU, the O-J transition amplitude increased from 39.1 to 58.6 % (Table 1), indicating that a blocked forward electron flow stimulated Q_A_^-^ accumulation, albeit to a much smaller extent.

In contrast to the other samples, DCMU does not significantly affect the OJIP transient in the PSII-I assembly intermediate. The O-J, J-I and I-P amplitudes show only minimal changes in the presence of DCMU (Fig. 3b, red trace). Notably, the O-J transition amplitude is increased by 5.4 % after DCMU addition, offering a similar yet less pronounced trend to that observed with other PSII complexes (Table 1). This observation supports our previous work, where the addition of DCMU did not significantly alter Q_A_^-^ reoxidation kinetics because of an immature Q_B_-binding pocket [1]. DCMU interacts with Ser264 and Phe265 of the D1 subunit [35]. In the PSII-I assembly intermediate, the auxiliary factor Psb28 distorts the Q_B_-binding pocket by reorientating Phe265 of the D1 subunit [1] (Fig. 1), which in the mature structure helps to orient the Q_B_ headgroup [6], ensuring its optimal position for forward electron transfer.

Interestingly, the control sample (Fig. 3a, blue trace) shows an OJIP fluorescence induction similar to traces observed in pre-illuminated pea leaf measurements. Pre-illumination ensured a reduced PQ-pool that increased the rate of Q_A_^-^ accumulation [33]. Similarly, the P-peak occurs due to Q_A_^-^ accumulation after over-reduction of the PQ-pool during the measurement [36]. The ratio between P and J-peak intensity was consequently increasing with increased reduction of the PQ-pool. In intact pea leaves, the J-peak intensity increases dramatically in measurements with pre-illuminated leaf material, indicating that this feature of the fluorescence induction profile can serve as an index of plastoquinone redox poise [33]. The fast accumulation of Q_A_^-^ is accelerated by adding urea-based pesticides, like DCMU, since they bind to the Q_B_-binding pocket, thereby preventing Q_B_ binding and forward electron transfer, mimicking an over-reduced PQ-pool. In contrast, measurements with isolated active PSII performed in the presence of the quinone analogue DCBQ (Fig. 3c, blue trace) had similar curve shapes to traces obtained with dark-adapted whole-cell measurements [19,31,33,37]. Most striking is the decreased O-J transition amplitude, dropping from 82.3 % to 59 % with the electron acceptor’s addition (Table 1), indicating a robust forward electron transfer to DCBQ and a slower accumulation of Q_A_^-^, more comparable to fluorescence induction traces measured *in-vivo*, demonstrating that this feature is extremely sensitive to the presence of electron acceptor molecules.

### 3.4. The Q_A_^-^ state is destabilized when the OEC is inactive

Figure 3a and b show that inactivation of the OEC causes an approximately 10-times faster onset of *Fm* than in the control PSII sample. In the PSII-I assembly intermediate, the combined effects of a missing OEC, distorted Q_B_-binding site and removed bicarbonate ligand of the non-heme iron lead to an increased charge dissipation rate through direct P_680_^+^/Q_A_^-^ recombination [1], caused by an increase in the redox potential of the Q_A_/Q_A_^-^ pair [13,30]. Furthermore, without the addition of DCMU, the O-J transition is lowest in the PSII-I and apo-PSII samples, namely 36.6 % and 39.1 %, respectively (Table 1). The lower O-J transition amplitudes indicate that inactivation of the OEC, either by Mn-depletion (apo-PSII) or by, to a lesser extent, removing the extrinsic PSII subunits PsbO, PsbU and PsbV (PSII(-OUV)) lead to a destabilization of the Q_A_^-^ state, visible by the lower O-J transition amplitudes and faster onset of the P-peak (Fig. 3a). This effect is most likely caused by a positive shift in Q_A_ redox potential [13,30], favoring charge dissipation through direct P_680_^+^/Q_A_^-^ charge recombination, as observed in the PSII-I assembly intermediate [1].

This trend is mirrored in the quantum yield efficiencies (*Fv/Fm*) of the different PSII complexes. Disruption of the OEC lowered the *Fv*/*Fm* ratio from 0.3 (*SD* = 0.004), as calculated with the control PSII sample, to 0.16 (*SD* = 0.002) with PSII(-OUV). Surprisingly, the quantum yield efficiency in the PSII-I (0.16, *SD* = 0.003) and apo-PSII sample (0.16, *SD* = 0.01) were equal to that measured with the OEC disrupted sample (Fig. 3f). This is in line with our previous observations, where the addition of DCMU did not affect the Q_A_ reoxidation kinetics of the PSII-I sample, indicating that in these samples charge dissipation manly occurs trough recombination and not forward electron transfer. The assembly intermediate namely showed increased direct P_680_^+^/Q_A_^-^ charge recombination through Q_A_^-^/Tyr_Z_ and Q_A_^-^ /Tyr_D_ on their respective timescales [1]. Therefore, in OEC disrupted samples the Q_A_^-^ is destabilized by an increase in recombination events, that lower the *Fv*/*Fm* values to a level observed with the PSII-I assembly intermediate.

### 3.5. OEC removal increases DCBQ binding affinity to levels observed with the PSII-I assembly intermediate

Fluorescence measurements of different PSII complexes versus an increasing DCBQ concentration, ranging from the millimolar to the nanomolar range, are shown in Figure 4. Titration of increasing concentrations of an electron acceptor to PSII decreased the fluorescence values measured at each acceptor concentration. Plastoquinone analogues, such as DCBQ, produce their fluorescence quenching effect by oxidizing Fe(II) through the Q_B_ binding pocket [38]. Oxidation occurs via an interaction with D1 His215 [35], a residue involved in coordinating the non-heme iron [6]. Fitting the fluorescence values to a specific binding with Hill slope model allowed us to determine the equilibrium disassociation constant (*Kd*) values and their associated hill slope (*h*) values.

**Fig. 4:**
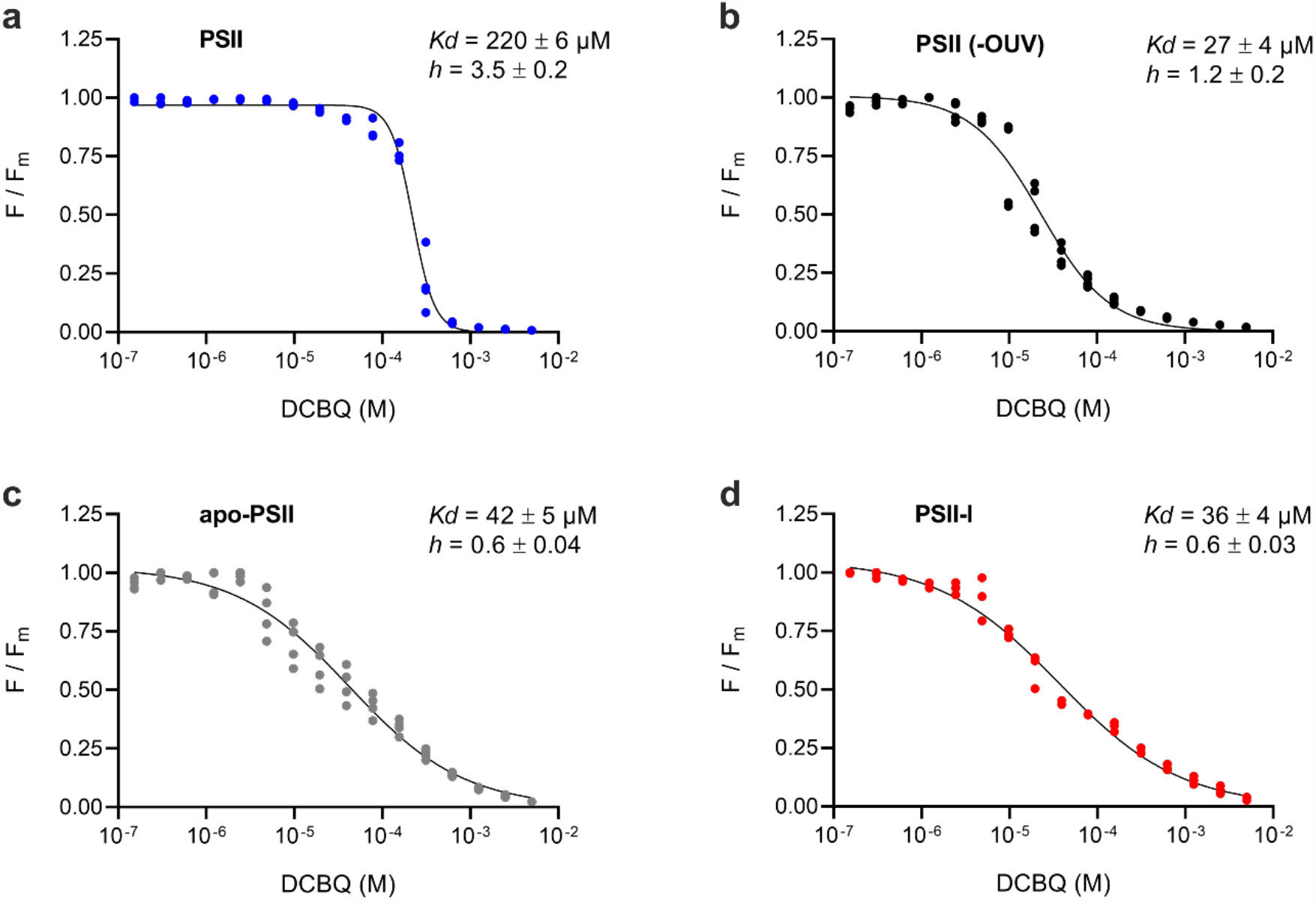
**Titration of DCBQ** to **(a)** PSII, **(b)** PSII(-OUV), **(c)** apo-PSII and **(d)** PSII-I. The dots depict normalised fluorescence values, of at least three independent measurements, at different DCMU concentrations, and the black line represents the nonlinear fit of the data. The equilibrium disassociation constant (*Kd*) and the hill slope value (*h*) are given in the individual graphs’ upper right corner with their corresponding standard error of mean (SEM) values.

Active dimeric PSII exhibited a *Kd* of 220 µM, with a Hill slope (*h*) of 3.5, indicating cooperative binding. Surprisingly, the inactivation of the OEC increased the affinity for DCBQ by approximately 10-fold (Fig. 4a, b and c). The shift in *Kd* is similar to the trend observed by Fagerlund et al., where the deletion of PsbT and subsequent accumulation of assembly modules produced an increased affinity towards 2,6-dimethoxybenzoquinone (DMBQ) [31]. Inactivation of the OEC by removal of the extrinsic subunits in the sample PSII(-OUV) decreased the *Kd* to 27 µM, with a 1.2 *h*-value. Similar results were obtained with the Mn-depleted apo-PSII sample. There the *Kd* was determined at 42 µM, with an 0.6 *h*-value. The results obtained with the apo-PSII sample were similar to the values measured with the PSII-I assembly intermediate. The affinity was slightly higher (*Kd* = 36 µM), with an 0.6 *h*-value indicating negative cooperative binding (Fig. 4d), that is characteristic for proteins with multiple binding sites with different affinities. The small *h*-value could also be a consequence of the inherent heterogeneity of the PSII-I sample [1].

## 4. Discussion

### 4.1. The availability of external electron acceptors has a significant impact on variable (OJIP) chlorophyll fluorescence induction traces

Variable OJIP fluorescence traces measured with isolated and purified PSII complexes show the characteristic O, J, I and P fluorescence induction features (Fig. 3a), although with distinct differences when compared to measurements performed with whole-cell material. The phase transition amplitudes without the addition of artificial electron acceptors (Table 1) are characterized by an increase in the O-J transition amplitude, compared with values published for whole-cell measurements [19,37]. When comparing values obtained with whole-leaf measurements and isolated thylakoid membranes in spinach, the O-J transition amplitude was the only parameter that increased [37], indicating that a lack of a PSII-external electron acceptor system induces faster accumulation of reduced immobile plastoquinone. A similar phenomenon was also observed with pre-illuminated pea leaves, where treatment with actinic light induces the closure of reaction centres by reducing the PQ-pool [33]. A comparable O-J transition amplitude increase can be achieved through the addition of DCMU [19,31–33], a urea-based herbicide that specifically blocks the Q_B_-binding pocket [35]. This effect is attributed to a faster accumulation of Q_A_^-^ facilitated by blocked forward electron transport, either by an over-reduced PQ-pool [33] or a blocked Q_B_-site, as is the case with DCMU [19,31,32,35].

We were able to observe this phenomenon with detergent solubilized PSII complexes. In our measurements, the absence of an external PQ-pool leads to a faster Q_A_^-^ accumulation, visible in a comparably high O-J transition amplitude (Fig. 3a and Table 1). The addition of DCMU amplified the same effect. There, the O-J transition accounts for almost all the fluorescence increase (Fig. 3b and Table 1). On the other hand, measurements performed in the presence of the quinone analogue DCBQ (Fig. 3c) produced curve shapes similar to traces obtained in dark-adapted whole-cell measurements [31,33]. These were characterized by a more pronounced intermediary I-inflection of the fluorescence induction curves (Fig. 3c). Similarly, adding an excess of oxidized decyl-plastoquinone (dPQ) to intact spinach chloroplasts induced a sharpening of the intermediary I-inflexion [37], mirroring our measurements with purified PSII complexes.

Our data obtained with solubilized and highly purified PSII complexes allowed us to isolate the effect of the PQ-pool on variable (OJIP) fluorescence induction traces in an *in vitro* experimental setup. In brief, the addition DCMU blocked forward electron transport and led to a faster accumulation of reduced immobile plastoquinone (Q_A_^-^). This effect is evident by the equalization of the P and J-peak intensities (Fig. 3b and Table 1), confirming prior observations made with whole-cell material [33]. Furthermore, we were able to show in an *in vitro* experimental setup that the addition of external PSII electron acceptors has the most significant impact on the intermediary inflexion-I of the OJIP transient (Fig. 3c), mirroring observations performed on intact spinach chloroplasts [37]. Thus, we can conclude that even small PQH_2_/PQ ratio changes can be indirectly observed through the OJIP fluorescence induction profile.

### 4.2. The functionality of the Mn-cluster determines variable fluorescence induction kinetics

The destabilization of the OEC (PSII(-OUV)) and Mn-depletion (apo-PSII) of PSII complexes resulted in the onset of a premature J-peak like feature (Fig. 3a) previously described as the so-called K-peak [26]. Removal of the extrinsic subunits in the PSII(-OUV) sample (Fig. 2a and b) resulted in a lower and slightly faster onset of the J-peak. The shift in J-peak onset and lowering of the O-J transition amplitude was even more pronounced in the Mn-depleted apo-PSII sample (Fig. 3a), with the decrease in amplitude mirrored in the calculated parameters. The PSII-I assembly intermediate exhibited the lowest O-J transition amplitude of 36.6 % (Table 1).

The PSII-I intermediate represents a snapshot of CP43 attachment to the RC47 complex. This is before the full maturation of the Q_B_-binding pocket and activation of the oxygen-evolving complex [1]. Several effects overlap in the PSII-I assembly intermediate to produce the low O-J transition amplitude observed (36.6 %). First, the absence of the OEC shifts the Q_A_/Q_A_^-^ redox potential by approximately +150 mV [30,39]. And second, a similar shift is caused by replacing bicarbonate as a ligand of the non-heme iron with format, with a reported + 70 mV increase in Q_A_/Q_A_^-^ redox potential [13]. Interestingly, this substitution shifts the characteristic EPR signal of the Q_A_^-^Fe^2+^/Q_B_^-^Fe^2+^ complexes from g = 1.9 [40] to g = 1.84, a value characteristic for the bacterial reaction centre [41]. In PSII-I the D1 Glu241 residue replaces bicarbonate as a ligand of the non-heme iron (Fig. 1). This shift in the position D1 Glu241 creates a similar coordination of the non-heme iron as is found in the bacterial reaction centre of *Rhodobacter sphaeroides* [1]. Therefore, it is conceivable that a similar shift in Q_A_/Q_A_^-^ redox potential also occurs in the PSII-I assembly intermediate.

Positive shifts in Q_A_/Q_A_^-^ redox potential favour direct and safe P_680_^+^/Q_A_^-^ charge recombination, producing a lower probability of ^3^P_680_ formation and lower ^1^O_2_ production rates [1,42]. Consequently, the Q_A_^-^ state becomes destabilised, and this is visible in the lower overall quantum yield efficiency (Fig. 3d, e and f) and lower J-peak intensity (Fig. 3a) that is mirrored in the lower calculated O-J transition amplitudes (Table 1) of the PSII-I complex. Although the direct Q_A_/Q_A_^-^ redox potential in the PSII-I intermediate was not determined, it showed a lower ^1^O_2_ production rate than was observed with inactivated PSII-I [1], indicating a shift to the high potential form of the Q_A_/Q_A_^-^ redox pair where ^1^O_2_ production is minimal [42]. Therefore, our data indicate that in isolated PSII samples, the O-J transition amplitude is directly connected to the Q_A_^-^ accumulation rate. In the absence of an external electron acceptor, the speed of accumulation is strongly affected by the redox potential of the Q_A_/Q_A_^-^ pair since it dictates the final fate of the excited electron [42]. The higher recombination rates observed in the PSII-I assembly intermediate [1], caused by the absence of the OEC [30,39] and bicarbonate ligand [13], led to slower Q_A_^-^ accumulation in the PSII-I intermediate (Fig. 3a). This mechanism is reflected in the lower O-J transition amplitude of the PSII-I assembly intermediate and, to a lesser extent, in the deactivated PSII complexes (Table 1). Therefore, we propose that the O-J transition amplitude can be used to assess the Q_A_/Q_A_^-^ redox potential in isolated PSII complexes. There the effect of the higher redox potential is isolated from the fluorescence quenching effect of the PQ-pool. Meaning that higher redox potentials will disfavour Q_A_^-^ accumulation, by an increased recombination rate [1,42] and consequently lower the J-peak amplitude.

### 4.3. Inactive PSII complexes have a higher affinity towards plastoquinone analogues

Docking simulations combined with FTIR data have revealed that Q_B_ binding involves forming a hydrogen bond between one of the plastoquinone carbonyl oxygen atoms and Ser264 of the D1 subunit. The remaining CO group forms a hydrogen bond with the backbone HN group of Phe265 and His215 of the D1 subunit. The latter interaction is absent when docking DCMU, a urea-based herbicide. On the other hand, the herbicide bromoxynil interacts only with D1 His215 through a hydrogen bond between its phenolic CO group and an imidazole NH group of histidine [35]. In the PSII-I assembly intermediate, the position of D1 Phe265 is altered by the action of the axillary factor Psb28 (Fig. 1). It forms an extended beta-hairpin structure with the D-E loop of the D1 subunit [1], a region involved in bicarbonate binding and the formation of the Q_B_-binding pocket [6]. Reorientation of the D1 Phe265 sidechain in PSII-I results in the variable (OJIP) fluorescence induction traces showing no change after the addition of DCMU (Fig. 3b). Surprisingly, despite the changes in the Q_B_-binding pocket, the quinone analogue DCBQ could still quench PSII-I fluorescence, thus indicating its functional binding to the complex. This is apparent in the drastic decrease of *Fv/Fm* induced by DCBQ (Fig. 3f) and the changes in OJIP fluorescence kinetics after its addition (Fig. 3c and Table 1). Together with the observation that DCMU does not affect Q_A_-reoxidation in PSII-I [1], these findings indicate that DCBQ has a binding mode apart from DCMU. We propose that DCBQ mainly interacts with the residues coordinating the none-heme iron, similar to the proposed binding model for bromoxynil [35], where it oxidizes Q_A_ through its interaction with D1 His215 within the Q_B_-binding pocket.

Our data suggest that the Q_A_ redox potential impacts quinone binding affinity. In samples where the OEC was either deactivated or entirely removed, the affinity of PSII towards DCBQ increased by approximately 10-fold. Interestingly, the PSII-I assembly intermediate exhibited a similarly high affinity as the artificially inactivated samples (Fig. 4b, c and d).

Removal of the OEC via hydroxylamine treatment [17] increases the Q_A_^-^/Q_A_ redox potential for approximately 150 mV [30,39]. Conversely, Mn-depletion produces only a relatively minor shift in the redox potential of the non-heme iron (+ 18 mV) [30]. This small change would not fully account for the drastic increase in DCBQ affinity we observed after OEC removal and in the PSII-I assembly intermediate. On the other hand, it was reported that this slight shift in non-heme iron redox potential is accompanied by a significant alteration of the *pK’*_*a*_ values of His215 and Glu241 of the D1 subunit. This is accompanied by slight structural changes to the D-E loop, that lead to a stronger interaction between the carbonyl oxygen of Q_B_ and the sidechain HN of His215 of the D1 subunit [30]. Our data support these suggestions, where the removal and even disruption of the OEC (Fig. 4b and c), caused an approximately 10-fold increase in binding affinity towards DCBQ, a plastoquinone analogue. Furthermore, the high affinity of PSII-I for DCBQ would indicate that D1 His215 is the driving force behind the binding event, the indication compounded by the similar affinities displayed by the two artificially inactivated PSII samples (PSII(-OUV) and apo-PSII). There, the integrity of the Q_B_-binding pocket is not altered by the Psb28-induced structural changes. The *Kd* observed in the PSII(-OUV) and apo-PSII samples were similar to the *Kd* calculated with the PSII-I assembly intermediate, them being 27, 42 and 36 µM, respectively (Fig. 4b, c and d). Therefore, we can conclude that a similar phenomenon occurs within the PSII-I assembly intermediate, as was reported for Mn-depleted PSII [30]. The positive shift in Q_A_/Q_A_^-^ redox potential, caused by the D-E loop’s reorientation by binding Psb28, and the absence of the OEC [1], strengthens the interaction between D1 His215 and DCBQ, which leads to an overall higher binding affinity. Interestingly, we were also able to observe this effect when titrating the plastoquinone analogue dPQ, with an apparent 10-fold increase in affinity when comparing active PSII (*Kd* = 698 ± 67 µM) with the PSII-I (*Kd* = 67 ± 10 µM) assembly intermediate.

A similar effect was already observed in a mutant strain of *Synechocystis* sp. PCC 6803, where a PsbT deletion caused an approximately two-fold higher affinity to the plastoquinone analogue DMBQ. The mutant was deficient in PSII assembly and accumulated PSII assembly modules, most notably RC47, with an accompanying decrease in mature dimeric PSII complexes. By plotting the *Fm* values of Q_A_ reoxidation measurements versus increasing DMBQ concentrations, they could determine a *Kd* of ∼210 µM for the wild-type cells and a *Kd* of ∼140 µM for the PsbT deletion strain [31]. The value calculated with the WT strain is verry similar to our values, obtained with highly pure active dimeric PSII complexes (*Kd* = 220 µM) (Fig. 4a). This observation presents an indication that the same mechanism that we observed *in vitro* might also occur *in vivo*. Therefore, we propose a mechanism by which D1 His215 remains permanently occupied by a plastoquinone molecule until activation of the water-splitting activity. This would produce an additional positive shift in the Q_A_/Q_A_^-^ redox potential by blocking forward electron flow, similar to the effect of DCMU on Q_A_ redox potential [43]. Consequently, this would favor charge recombination via the direct P_680_^+^/Q_A_^-^ pathway that disfavors the production of ^1^O_2_ by bypassing the triplet chlorophyll state [42] and protecting PSII during biogenesis.

## 5. Conclusion

Our publication presents an *in vitro* variable (OJIP) chlorophyll fluorescence induction characterization with isolated and highly pure PSII complexes. We were able to show that the O-J transition is proportional to the PQH_2_/PQ ratio of the cell and that the same feature can serve as a gauge of Q_A_/Q_A_^-^ redox potential in isolated PSII complexes removed from the quenching effect of the PQ-pool. Furthermore, titration of a plastoquinone analogue to the PSII-I assembly intermediate gave us insight into the functionality of the Q_B_-binding pocket during PSII biogenesis. We showed that an increase in Q_A_/Q_A_^-^ redox potential is accompanied by an increase in the affinity towards plastoquinone analogues. Such an increase in affinity might serve a photoprotective role by adding a positive shift to the Q_A_/Q_A_^-^ during PSII assembly to minimize singlet oxygen production, a harmful reactive oxygen species.

## Acknowledgements

We thank M. Völkel for excellent technical assistance. Financial support was provided by the Deutsche Forschungsgemeinschaft (DFG) Research Unit FOR2092 (836/3-2 to M.M.N.).

